# A high-throughput fluidic chip for rapid phenotypic antibiotic susceptibility testing

**DOI:** 10.1101/647909

**Authors:** Pikkei Wistrand-Yuen, Christer Malmberg, Nikos Fatsis-Kavalopoulos, Moritz Lübke, Thomas Tängdén, Johan Kreuger

**Author notes:** PWY and CM have contributed equally to this manuscript.

## Abstract

Many patients with severe infections receive inappropriate empirical treatment and rapid detection of bacterial antibiotic susceptibility can in this context improve clinical outcome and reduce mortality. We have to this end developed a high-throughput fluidic chip for rapid phenotypic antibiotic susceptibility testing of bacteria. A total of 21 clinical isolates of *Escherichia coli, Klebsiella pneumoniae* and *Staphylococcus aureus* were acquired from the EUCAST Development Laboratory and tested against amikacin, ceftazidime and meropenem (Gramnegative bacteria) or gentamicin, ofloxacin and tetracycline (Gram-positive bacteria). The bacterial samples were mixed with agarose and loaded in 8 separate growth chambers in the fluidic chip. The chip was thereafter connected to a reservoir lid containing different antibiotics and a pump used to draw growth media with or without antibiotics into the chip for generation of diffusion-limited antibiotic gradients in the growth chambers. Bacterial microcolony growth was monitored using darkfield time-lapse microscopy and quantified using a cluster image analysis algorithm. Minimum inhibitory concentration (MIC) values were automatically obtained by tracking the growth rates of individual microcolonies in different regions of antibiotic gradients. Stable MIC values were obtained within 2-4 hours and the results showed categorical agreement to reference MIC values as determined with broth microdilution in 86% of the cases.

**Importance:** Prompt and effective antimicrobial therapy is crucial for the management of patients with severe bacterial infections but is becoming increasingly difficult to provide due to emerging antibiotic resistance. The traditional methods for antibiotic susceptibility testing (AST) used in most clinical laboratories are reliable but slow with turnaround times of 2-3 days, which necessitates the use of empirical therapy with broad-spectrum antibiotics. There is a great need for fast and reliable AST methods that enable start of targeted treatment within a few hours to improve patient outcome and reduce overuse of broad-spectrum antibiotics. The high-throughput fluidic chip for phenotypic AST described in the present study enables data on antimicrobial resistance within 2-4 hours allowing for an early initiation of appropriate antibiotic therapy.

## Introduction

Antibiotics are among the most successful drugs developed and has helped to drastically reduce mortality and morbidity from common bacterial infections such as pneumonia, urinary tract infections and bloodstream infections over the last century (1). Unfortunately, many antibiotics are becoming less effective due to the selection and spread of antibiotic-resistant bacteria (2)(3)(4), resulting from antibiotic overuse (5). The increasing resistance to standard antibiotic therapy increases the risk of inappropriate empirical antibiotic therapy, treatment failure and mortality in critically ill patients (6). For example, in southern and eastern Europe carbapenem resistance in Enterobacteriales is approaching 50% incidence (7)(8). Thus, new rapid and robust methods for antibiotic susceptibility testing (AST) that can deliver data on bacterial antibiotic resistance within hours instead of days are much needed. Such methods will make it possible to prescribe targeted antibiotic therapy, which will improve clinical outcome and reduce unnecessary use of broad-spectrum antibiotics, thereby minimizing the risk of adverse events and resistance development (9)(10)(11).

The traditional methods for AST currently used in most clinical microbiology laboratories are reliable and inexpensive but comparatively slow with turnaround times (TATs) commonly of 2-3 days (12). Progress has recently been made to decrease TATs, in part through automation of existing methods but also as a result of the implementation of new technologies (12). For example, recent developments of the rapid disc diffusion test with new breakpoints (4, 6 and 8 hour readouts) developed by EUCAST allow for TATs of around 24 hours, although not for all samples (13). Importantly, studies show that in severe infections such as bloodstream infections with septic shock and bacterial meningitis, every hour of delayed appropriate antibiotic treatment increases mortality (11).

Many new rapid AST methods under development are phenotypic tests, as genotypic tests for selected resistance markers currently cannot reliably predict antibiotic susceptibility due to the complexity of the bacterial resistome (14)(9)(12). The latest developments in resistance screening methods include mass spectrometry to identify resistance-associated proteins and degradation products of antibiotics, PCR-based techniques to detect resistance markers, RNA microarrays, and whole genome sequencing together with AI-based deep learning tools (14)(15). Novel microfluidic systems for phenotypic antibiotic susceptibility testing are also being developed by several research labs (16).

We have previously designed and evaluated a rapid phenotypic antibiotic susceptibility test where bacteria are mixed with agarose to form a gel within a growth chamber inside a microfluidic system, and subjected to a gradient of an antibiotic to enable MIC determination (17). The aim of the current study was to develop a high-throughput system (based on our previous design) capable of simultaneously analyzing 8 samples in one chip and with the possibility of testing several antibiotics in parallel. To accomplish this, we developed a high-throughput microfluidic chip using 3D-printed molds for PDMS casting, together with a 3D-printed chip holder with an integrated reservoir lid. Bacterial microcolony growth was monitored using darkfield time-lapse microscopy and microcolony growth quantified using a cluster analysis algorithm. MIC values were obtained by analysis of bacterial growth rates in different regions of the antibiotic gradients. Clinical isolates of *Escherichia coli, Klebsiella pneumoniae* and *Staphylococcus aureus* with different antibiotic susceptibility profiles were tested against 6 commonly used antibiotics and stable MIC values obtained within 2-4 hours.

## Results

### Design and manufacture of the fluidic system

The goal of the present study was to develop a high-throughput fluidic system that allows for parallel and automatic analyses of 8 independent samples to obtain MIC values for different antibiotics. The construction of the high-throughput system required several new technical solutions. In the fluidic chip, each growth chamber - flanked by two fluidic channels used for delivery of growth media and antibiotics - was designed to feature a loading port that was used to inject bacteria mixed with agarose (Figure 1A-C). The growth chamber was designed to hold a volume of 5 μl. Loading of the bacteria-agarose mix filled up the growth chamber all the way to the edges of the flanking fluidic channels, which were designed to be slightly enlarged at the location of the growth chamber to tolerate slight variations in agarose gel volume (e.g. overloading) without compromising fluid flow. Consistent with our previous study (17), there was no fluid flow through the gel–filled growth chamber. Instead, molecules travel through the growth chamber between the two flanking flow channels by diffusion. Thus, diffusion-limited antibiotic gradients can be established through the agarose gel by the addition of an antibiotic to one of the two flow channels that flank a growth chamber (Figure 1A). Importantly, the constant fluid flow on both sides of the growth chamber establishes a stable source-sink system. MIC values are determined by analyzing bacteria microcolony growth in the different regions of the established antibiotic gradient that correspond to distinct antibiotic concentrations (Figure 1A).

**Figure 1.**
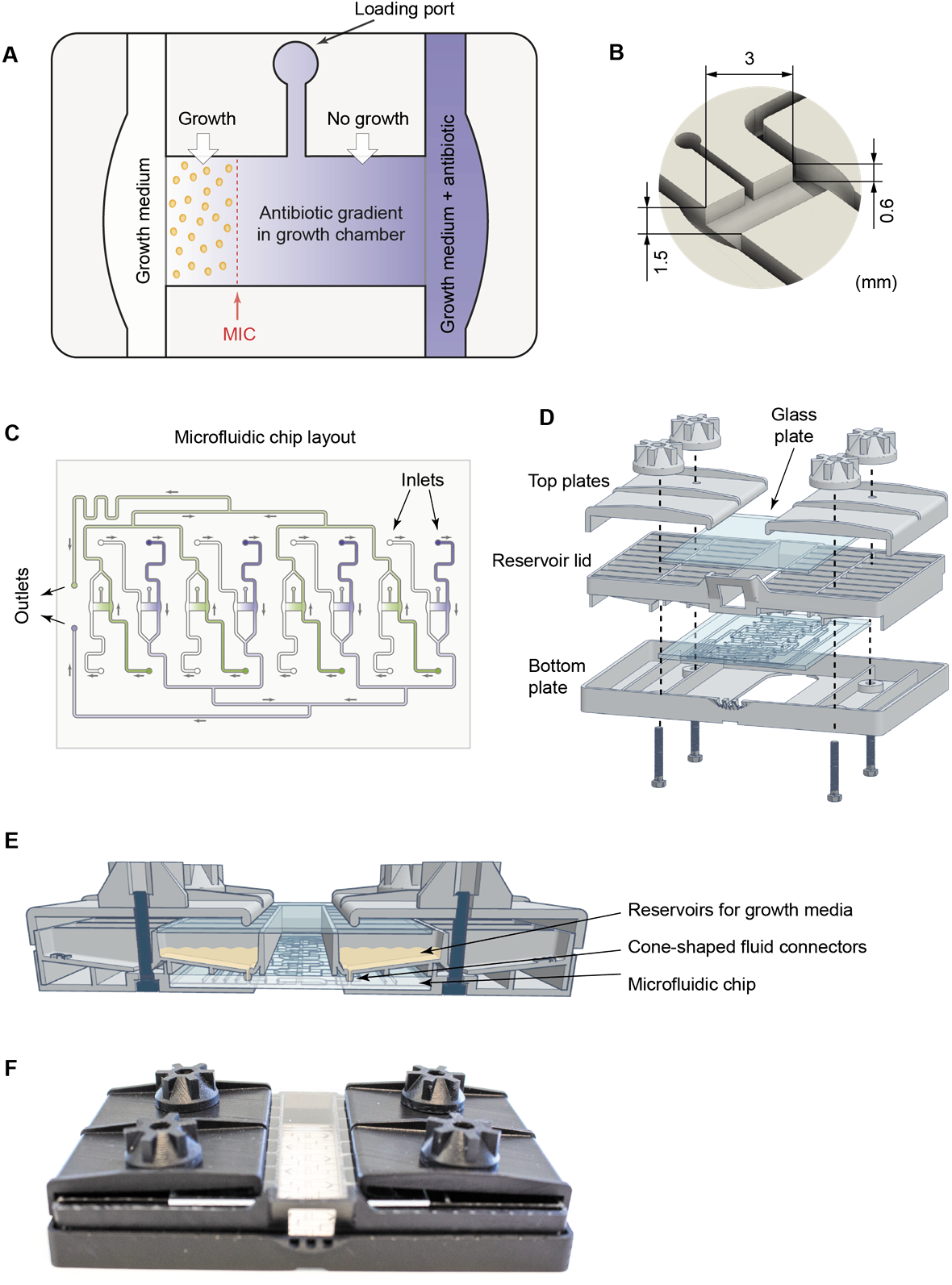
Overview of the fluidic system. (A) Illustration showing a growth chamber and a loading port used to inject the bacteria-agarose mix. The growth chamber is flanked by two flow channels. Antibiotic gradients in the growth chamber are created by diffusion from the source channel (blue) containing growth medium as well as a high concentration of the antibiotic to the sink channel (white) containing only growth medium. Bacteria present in the growth chamber are exposed to the diffusion-limited antibiotic gradient, and the MIC value will correspond to the lowest concentration at which the bacteria do not grow to form microcolonies (shown in this example as a red dotted line). (B) A close-up view of a growth chamber with dimensions indicated. (C) Overview of the microfluidic chip holding 8 growth chambers. The fluidic outlets that are connected to a pump are indicated, as well as the inlets (16 in total) that are connected to the reservoir lid. (D) Drawing showing how the bottom plate, the fluidic chip, the reservoir lid, and the top plates are assembled. (E) A cartoon showing the assembled system from the side highlighting the connections between the reservoir lid and the fluidic chip via fluid connectors. (F) A photograph of the assembled system.

In order to optimally position the 8 growth chambers together with their associated flow channels, fluidic inlets and outlets, the growth chambers were divided into two groups that within each group had the same orientation and shared a common outlet channel (Figure 1C). The chip was generated by casting PDMS into 3D-printed molds coding for the fluidic channels and growth chambers. The resultant PDMS chip was then removed from the mold where after the PDMS chip was bonded to a glass plate to create a closed fluidic system (see materials and methods for a detailed description). The fluidic chip was connected to a 3D-printed reservoir lid that was part of a larger chip holder (Figure 1D-F). The lid had 16 medium reservoirs with small cone-shaped fluid connectors in the bottom, which could be pressed into the chip inlets (Figure 1E), to enable delivery of growth medium with or without antibiotics from the reservoirs to the different fluidic channels of the system. The reservoir lid was held together with the microfluidic chip, a 3D-printed bottom plate and two top plates (including a glass plate positioned above the medium reservoirs to prevent evaporation) by M3 screws passing through the bottom plate, reservoir lid and top plates to ensure tight fluid connections between the lid reservoirs and the chip (Figure 1D-F).

### System validation

A fluorescein dye was used as a model molecule for antibiotic diffusion, to visualize and validate the formation of antibiotic gradients in the growth chambers. The system was operated using a syringe pump (see the Materials and methods section for details) and quantifications of the gradients performed after 2 hours, when the fluorescein gradients were considered to be fully established. Notably, the homogeneity of the formed gradients in the growth chambers was clearly affected by the loading port channel, which acted as a sink (Figure 2A). The half of the growth chamber closest to the loading port was therefore later to be excluded from the image analysis of bacteria growth. The gradients formed in the other half of the growth chamber (i.e. distal to the loading port) were shown to be linear (R^2^=0.988) and consistent between different chambers and chips (Figure 2).

**Figure 2.**
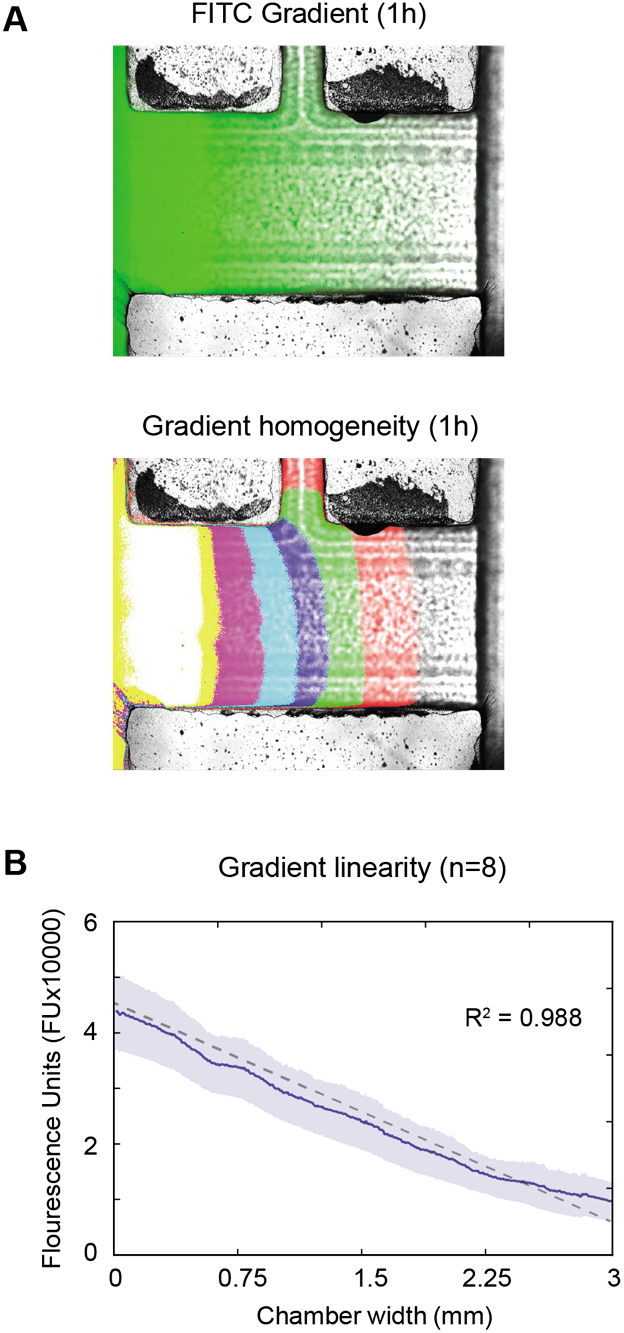
Gradient characterization. (A) Gradient formation in the growth chambers was validated using fluorescein. (B) The average fluorescein gradient (solid line, 95% confidence interval indicated by blue area) and the linear least squares fit (dotted line) of the data (*n* = 24).

### Data acquisition and calculation of MIC values

An automated dark-field microscope was used to perform time-lapse imaging of every growth chamber in the fluidic chip, and images from the different growth chambers were acquired every 10 minutes. Figure 3 shows an example where *E. coli* were subjected to a gradient of amikacin, to illustrate how the data was acquired and handled in order to obtain MIC values. An image analysis algorithm was developed to extract data from the series of images recorded from the growth chambers for identification of bacterial microcolonies, tracking their growth over time (Figure 3A, B) to automatically determine MIC values (Figure 3C-F). Briefly, images of the growth chambers were first corrected for mechanical drift. The images were thereafter cropped to include only the part of the growth chambers where the antibiotic gradients were linear, followed by subtraction of background signal. To be able to quantify growth, each image was binarized so that every colony was represented by a cluster of positive pixels, and the size of the cluster was followed and compared over time. Bacteria microcolonies unaffected by the local antibiotic concentration initially showed a linear growth pattern, whereas non-growing colonies exhibited a constant size throughout the experiment (Figure 3C-D). Colony growth was next expressed as a function of antibiotic concentration and a Gaussian fit used to calculate a MIC value at every time point. A final MIC value was reported once the corresponding positive growth chamber had flagged positive and if the calculated MIC value remained stable (+/− 5%) for 30 minutes (Figure 3 E-F) or alternatively reported as the value obtained at the very end of the run (at *t* = 300 min).

**Figure 3.**
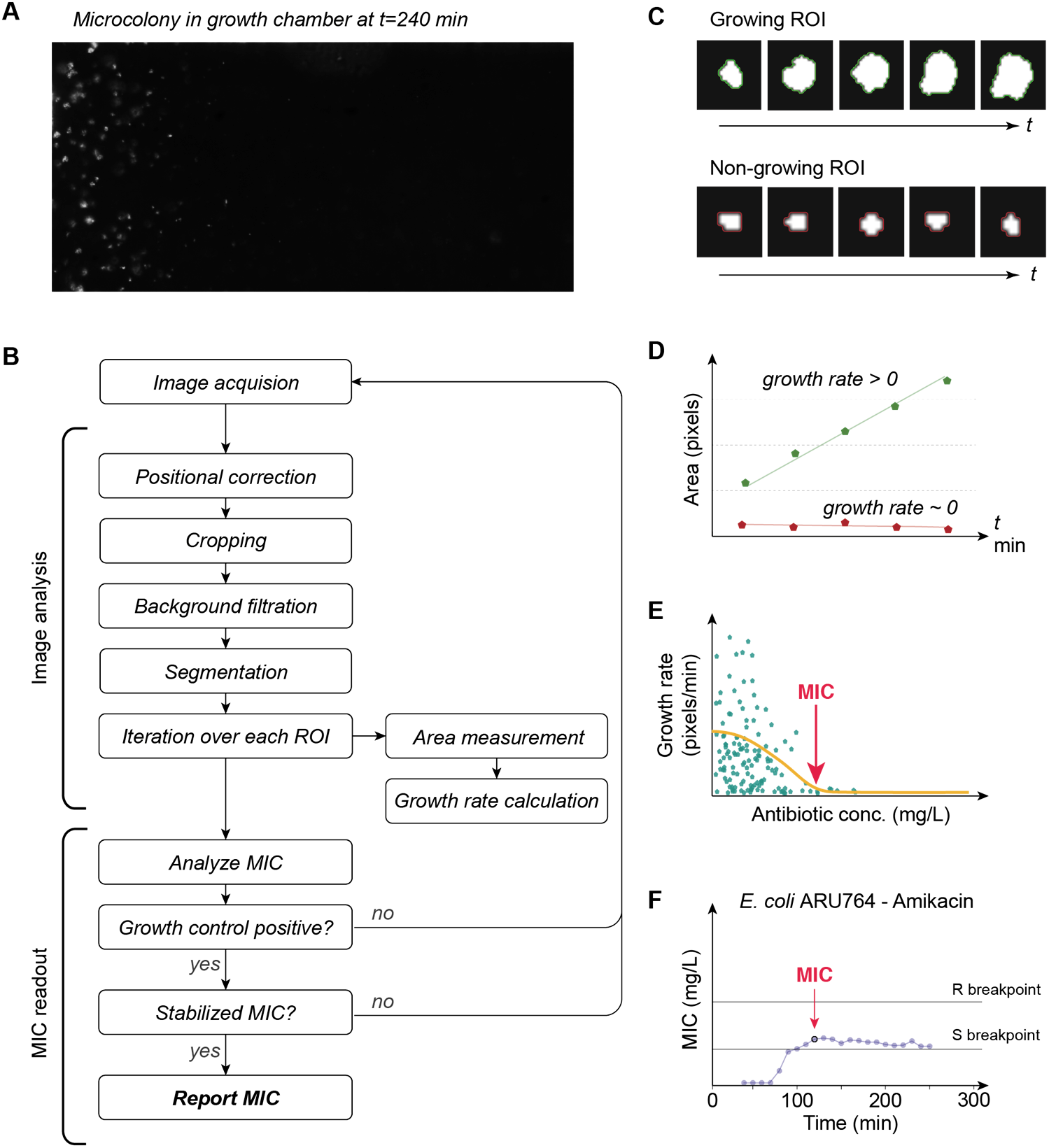
Data analysis. (A) Example showing *E. coli* grown in a gradient of amikacin. (B) Flow chart summarizing key actions of the image analysis algorithm. (C, D) Bacteria microcolonies that are exposed to antibiotic concentrations below MIC (growing ROI) keep expanding over time (green line in D), whereas colonies exposed to antibiotic concentrations at or above MIC (non-growing ROI) show no growth (red line in panel D). (E) Colony growth expressed as a function of antibiotic concentration: MIC values were calculated for every time point. (F) A final MIC value was reported if it had remained stable (+/− 5%) for 30 minutes.

### Rapid antibiotic susceptibility testing

After testing of the fluidic system and optimization of the accompanying analysis algorithm, we performed rapid AST of 21 clinical isolates: 11 Gram-negative (G-) strains (6 *E. coli* and 5 *K. pneumonia*) and 10 Gram-positive (G+) strains (*S. aureus*). Every strain was tested 4 times against three antibiotics: amikacin, ceftazidime and meropenem for the G− strains, and gentamicin, ofloxacin and tetracycline for the G+ strains. A positive control growth chamber containing bacteria but no antibiotics was included for each replicate (Supplementary Material S1). All MIC values measured in the fluidic chip were compared to reference MICs generated with the broth microdilution (BMD) method (Table 1).

**Table 1.** Comparison of MIC values obtained using BMD or the here developed fluidic chip assay.

**A**
| MIC (mg/L) using reference method (BMD) or the fluidic chip (FC) for G- strains |  |  |  |  |  |  |  |
| --- | --- | --- | --- | --- | --- | --- | --- |
| Strain nr. | Species | Amikacin |  | Ceftazidime |  | Meropenem |  |
|  |  | BMD | FC | BMD | FC | BMD | FC |
| ARU764 | *E. coli* | 16 | 16 | >16 | ≥16 | >16 | 4 |
| ARU754 | *E. coli* | >64 | ≥64 | >16 | ≥16 | 16 | ≤1 |
| ARU755 | *E. coli* | 4 | ≤2 | ≤0.5 | ≤0.5 | ≤1 | ≤1 |
| ARU756 | *E. coli* | ≤2 | ≤2 | 8 | ≥16 | ≤1 | ≤1 |
| ARU757 | *E. coli* | 4 | ≤2 | 1 | ≤0.5 | ≤1 | ≤1 |
| ARU758 | *E. coli* | 8 | ≤2 | ≤0.5 | 2 | ≤1 | ≤1 |
| ARU759 | *K. pneumoniae* | 4 | ≤2 | >16 | 8 | 4 | ≤1 |
| ARU760 | *K. pneumoniae* | 4 | ≤2 | >16 | ≥16 | 2 | ≤1 |
| ARU761 | *K. pneumoniae* | ≤2 | ≤2 | ≤0.5 | ≤0.5 | ≤1 | ≤1 |
| ARU762 | *K. pneumoniae* | ≤2 | ≤2 | ≤0.5 | ≤0.5 | ≤1 | ≤1 |
| ARU763 | *K. pneumoniae* | ≤2 | ≤2 | 1 | 8 | ≤1 | ≤1 |

**B**
| MIC (mg/L) using reference method (BMD) or the fluidic chip (FC) for G+ strains |  |  |  |  |  |  |  |
| --- | --- | --- | --- | --- | --- | --- | --- |
| Strain nr. | Species | Gentamicin |  | Ofloxacin |  | Tetracycline |  |
|  |  | BMD | FC | BMD | FC | BMD | FC |
| ARU795 | *S. aureus* | 0.25 | ≤0.125 | 1 | 1 | 0.5 | ≤0.25 |
| ARU796 | *S. aureus* | 0.25 | ≤0.125 | 2 | 2 | >8 | 4 |
| ARU797 | *S. aureus* | 0.5 | ≤0.125 | 0.5 | 0.5 | 0.5 | ≤0.25 |
| ARU798 | *S. aureus* | 0.5 | ≤0.125 | 0.5 | 1 | 1 | ≤0.25 |
| ARU799 | *S. aureus* | 0.5 | ≤0.125 | 2 | 2 | >8 | ≥8 |
| ARU800 | *S. aureus* | 0.5 | 0.25 | >4 | ≥4 | 2 | 0.5 |
| ARU801 | *S. aureus* | 1 | ≤0.125 | 0.5 | 0.5 | 0.5 | ≤0.25 |
| ARU802 | *S. aureus* | 1 | 0.25 | 1 | 1 | ≤0.25 | ≤0.25 |
| ARU803 | *S. aureus* | 2 | 1 | 0.5 | ≤0.125 | ≤0.25 | ≤0.25 |
| ARU804 | *S. aureus* | 2 | ≤0.125 | >4 | ≥4 | 4 | ≤0.25 |

Figure 4 shows MIC calculations over time for three representative strains (data for all strains is shown in Supplementary Material S1). The plotted line in each graph represents the mean MIC of 4 replicates and the blue area correspond to the standard deviation, whereas the dotted horizontal line signifies the MIC value obtained from reference measurements using BMD and the grey area corresponds to the accepted method variation of BMD. In the examples of *E. coli* representative strain (ARU756) and *K. pneumoniae* representative strain (ARU762), both amikacin and meropenem MICs were below the limit of quantification (LOQ) throughout the experiment (Figure 4), meanwhile when exposed to ceftazidime the strains initially appeared to grow unhindered during the first 120 minutes followed by a drop in MIC value. In *E. coli* ARU756, a ceftazidime-resistant strain, the MIC only decreased slightly and the resulting MIC was within range as compared with the BMD MIC. For *K. pneumoniae* ARU762, a ceftazidime-susceptible strain, a more apparent drop in MIC from resistant to susceptible below LOQ was observed. Similar “falsepositive” peaks were evident in several strains sensitive to ceftazidime and were likely due to phenotypic responses resulting in morphological changes such as filamentation. Filamentation is a well-known phenotypic effect of PBP3-targeting beta-lactams like ceftazidime (18)(19).

**Figure 4.**
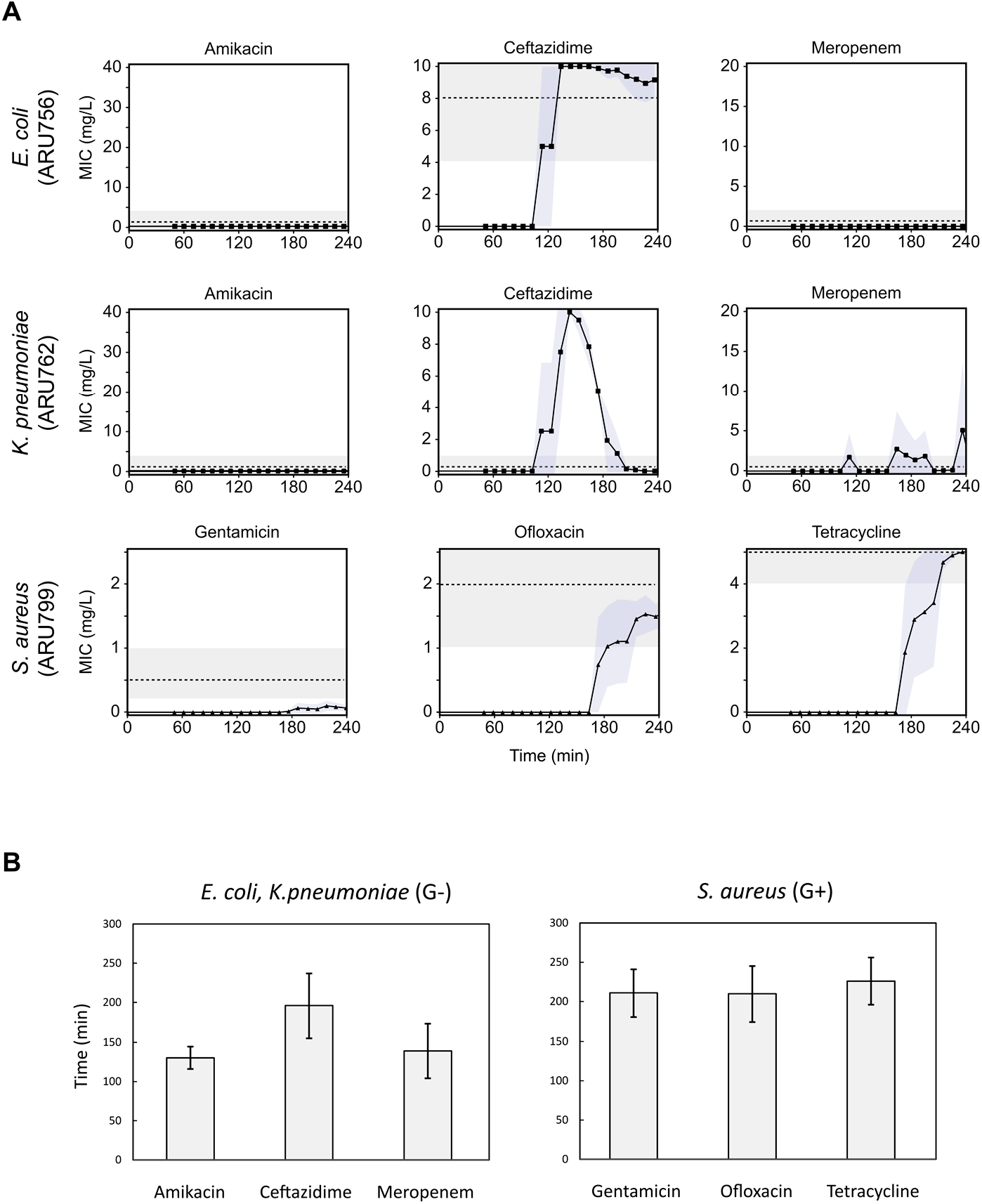
Examples of MIC values. (A) Data from representative strains of *E. coli, K. pneumoniae* and *S. aureus* detailing the MIC signal response over time for amikacin, ceftazidime and meropenem (*E. coli, K. pneumoniae*) and gentamicin, ofloxacin and tetracycline (*S. aureus*) respectively. Blue fields = SD, n = 4. The dotted lines correspond to the reference BMD MIC values, where visible in the tested concentration range. Grey fields correspond to acceptable variation of BMD, +− 1 log2 dilution. (B) Average read-out times until a stable MIC value after start of analysis for G− (left) and G+ (right) bacteria, error bars = SD.

For the antibiotics tested against G− isolates, 100% categorical agreement was observed for amikacin, 82% for ceftazidime and 74% for meropenem (Table 2). For the antibiotics tested against G+ isolates the categorical agreement was 80%, 100% and 80% for gentamicin, ofloxacin and tetracycline, respectively. Total categorical agreement between the new fluidic chip and BMD was 85.6%, with a range from 67-100%, and very major error (VME, when resistant strains are misclassified as susceptible), major error (ME, when susceptible strains are misclassified as resistant) and minor error (MiE, where resistant or susceptible strains are misclassified as sensitive with increased exposure, or sensitive with increased exposure strains misclassified as susceptible or resistant) were 6.3, 1.6 and 6.3% respectively (Table 2).

**Table 2.** Categorical agreement between BMD tests and the here described fluidic chip assay. Categorical agreement was scored according to the S-I-R classification (where S = sensitive; I = sensitive with Increased exposure, and R = resistant). Total categorical agreement between the MIC values obtained by the new fluidic chip and reference BMD method was 85.6%, with a range from 67-100%, and very major (VME), major (ME) and minor error (MiE) rates were 6.3, 1.6 and 6.3% respectively.

| Species | Antibiotic | No. of strains categorized in S, I, R according to the fluidic chip assay (BMD categorization shown within parenthesis<sup>a</sup>) |  |  | No. of CA (%) | No. of ME (%) | No. of VME (%) | No. of MiE (%) |
| --- | --- | --- | --- | --- | --- | --- | --- | --- |
|  |  | S | I | R |  |  |  |  |
| *E. coli* ($n = 6$) | AMI | 4 (4) | 1 (1) | 1 (1) | 6 (100) | 0 (0) | 0 (0) | 0 (0) |
|  | CAZ | 2 (3) | 1 (0) | 3 (3) | 5 (83) | 0 (0) | 0 (0) | 1 (17) |
|  | MER | 5 (4) | 1 (0) | 0 (2) | 4 (67) | 0 (0) | 1 (17) | 1 (17) |
| *K. pneumoniae* ($n = 5$) | AMI | 5 (5) | 0 (0) | 0 (0) | 5 (100) | 0 (0) | 0 (0) | 0 (0) |
|  | CAZ | 2 (3) | 0 (0) | 3 (2) | 4 (80) | 1 (20) | 0 (0) | 0 (0) |
|  | MER | 5 (4) | 0 (1) | 0 (0) | 4 (80) | 0 (0) | 0 (0) | 1 (20) |
| *S. aureus* ($n = 10$) | GEN | 10 (8) | 0 (0) | 0 (2) | 8 (80) | 0 (0) | 2 (20) | 0 (0) |
|  | OFL | 6 (6) | 0 (0) | 4 (4) | 10 (100) | 0 (0) | 0 (0) | 0 (0) |
|  | TET | 8 (6) | 0 (1) | 2 (3) | 8 (80) | 0 (0) | 1 (10) | 1 (10) |
| Total |  | 47 (43) | 3 (3) | 13 (17) | 54 (85.6) | 1 (1.6) | 4 (6.3) | 4 (6.3) |

<sup>a</sup> According to EUCAST clinical breakpoints table version 9.0
AMI, amikacin; CAZ, ceftazidime; MER, meropenem; GEN, gentamicin; OFL, ofloxacin; TET, tetracycline

Times-to-result were on average 155 min (SD: 43 min) for G− isolates (*n* = 132) and 216 min (SD: 32 min) for G+ isolates (*n* = 120), consistent with a more rapid growth rate of G− strains (Figure 4B).

## Discussion

The method presented here is able to rapidly provide antibiotic susceptibility data for up to 8 samples simultaneously, within 2-4 hours after the start of measurement in 89% of the tests, and within 5 hours for 100% of the tests. This is significantly shorter than traditional diagnostics such as disc diffusion and BMD which requires 18-24 hours. The method provides high information density per test chamber due to the analysis of bacteria growing in antibiotic gradients and has higher resolution than reference BMD tests and similar methods, which usually are based on discrete concentration steps dispersed over a bi-logarithmic scale. When comparing the results obtained using the here described fluidic chip to BMD, the poor essential agreement for some antibiotics can be explained by this difference, as isolates which have MIC-values in the lower concentration ranges cannot be well resolved at the lower end of the antibiotic gradient. The comparatively narrow antibiotic concentration range is the main disadvantage of the novel method, and inevitably many strains will have a MIC outside of the linear interval. These strains are then reported as MIC below LOQ or above max concentration, similarly to disc diffusion test which also has a narrower concentration range than BMD. Notably for the presented method, the linear gradient was positioned to include the S, I and R breakpoints for each tested antibiotic, as well as one log-2 dilution step above the R and below the S breakpoint, which is the range that arguably is of highest clinical importance. The narrow range provides an explanation as to why the categorical agreement to BMD was higher than the essential agreement, which is in line with existing comparative studies on AST methods (20),(21). On the other hand, the main advantage of using a linear gradient is that the resolution is higher in the clinically important range, with significantly lower variation than BMD, as seen in Figure 4.

The performance of the current method is clearly depending on i) isolate characteristics such as growth rate, ii) the type of antibiotic, as well as iii) the type of resistance. Firstly, phenotypic AST is dependent on measuring the growth, or absence thereof, of the bacteria. Therefore, slower growing bacteria will need relatively longer measurement times, no matter the technology used to measure growth. As an example, one of the *S. aureus* isolates included in this study (ARU803) grew very slowly and was the only strain where a MIC value could not be detected within 4 hours for any of the tested antibiotics. Secondly, the type of antibiotic affects the results, since the specific mode of action of an antibiotic will result in different lag-times before measurable phenotypic effects can be detected. For example, ceftazidime had a long lag-phase of ~2 hours before apparent growth stopped in susceptible bacteria (Figure 4). Consequently, the analysis algorithm tended to call the ceftazidime MIC values too early and therefore to classify all isolates as resistant, even those that were sensitive. However, by using information on growth patterns provided by the antibiotic-free growth control chamber, and by delaying the time until read-out for ceftazidime, this problem could be avoided. Thirdly, the type of resistance can also affect the results, as is likely the case for meropenem where growth for none of the two included resistant isolates were detected within 4 hours. However, when these samples were run for a longer period of time, bacterial growth was detectable (data not shown).

All rapid phenotypic AST methods will face the above discussed biological challenges to some extent, and whether a specific method will turn out to be viable will hinge on if the benefit of the rapid read-outs for a subgroup of patient isolates outweigh the increase in false or unclear readouts as well as the increased cost (9). Thus, rapid phenotypic AST methods will likely never completely replace traditional AST testing, but instead serve as an important addition to be used for example for critically ill patients (12).

In recent years, several microfluidics-based phenotypic rapid AST methods have been described in the literature, based on either direct or indirect growth markers (22). However, these methods often suffer from complicated sample preparation and loading, require pure cultures and/or specific strains (22), depend on complicated and expensive analysis instruments (22) or indirect markers (23), lack population scale data (24), or have limited output spectrum of only a single antibiotic concentration (25). Further, many emerging methods often measure effect on bacterial growth at a single antibiotic concentration set at the S or R breakpoint, thus limiting the resolution and clinical utility; or need multiple parallel wells for discrete antibiotic concentrations which increases the complexity and cost. In contrast, the fluidic system presented in this study is easy to use, robust, and provides growth data on bacteria growing in a continuum of antibiotic concentrations, thus avoiding these common drawbacks. We conclude that the method presented in this study has the potential to provide very rapid antibiotic susceptibility results of up to 8 samples and antibiotics per chip, which potentially allows an earlier switch to appropriate targeted therapy with narrow-spectrum antibiotics, improved clinical outcome as well as a lower risk of side effects and resistance development.

## Material and methods

### Construction of the microfluidic system

The molds for PDMS chip casting, as well as the reservoir lid and the bottom plate, were generated using a Form 2 3D-printer (Formlabs, Somerville, MA, USA) and printed horizontally using Black Resin (Formlabs) and 50 μm thick layers. The microfluidic chips were generated by casting PDMS (Sylgard 184, Sigma-Aldrich) into 3D-printed molds. The resultant PDMS chips were bonded to glass plates (62 x 82 x 1 mm, Marienfeld Superior, Paul Marienfeld GmbH & Co, Germany) with a calibrated corona treater (BD20-AC, ETP, Chicago, IL, USA) essentially as previously described (26).

### Validation of gradient formation

Fluid flows were created in the fluidic chip using a syringe pump (Chemyx Fusion 200, Chemyx, Stafford, TX, USA) set in withdrawal mode and flows of 1 μL/min were for all experiments generated in the chip flow channels. The fluorescent dye Fluorescein (#46955, Sigma Aldrich) was used to visualize gradient formation, and images of the formed gradients captured using a Axiovert 200M fluorescent microscope (Carl Zeiss, Jena, Germany). The quantification of the gradient was done using ImageJ1.8.0 (NIH, Bethesda, USA) and the values normalized to the highest signal in the source channel. Linear regression was carried out using linear square fitting, and standard deviations calculated using GraphPad Prism version 6 (GraphPad Software, San Dieago, CA, USA).

### Bacterial isolates

All bacteria strains were provided by the EUCAST Development Laboratory in Växjö, Sweden. Isolates of *E. coli* (n = 6), *K. pneumoniae* (n = 5) and *S. aureus* (n = 10) with varying susceptibility against the tested antibiotics were used in this study (see Table 1). All strains were cultivated using cation-adjusted Müller-Hinton (MH-II) broth and MH-II agar.

### Antibiotics

Antibiotics were purchased from Sigma Aldrich; ceftazidime and ofloxacin were the European Pharmacopeia standards and the meropenem used was the US Pharmacopeia standard. All antibiotics were first prepared as 20 x stock solutions and stored at −80 °C until use. The antibiotic concentrations used in the experiments in the source solutions for gradient generation were 2.5-fold higher than the clinical breakpoints for resistance for the tested antibiotic against the tested specimen in each specific case according to the EUCAST Clinical Breakpoints Table 8.0. Amikacin, ceftazidime and meropenem were tested against *E. coli* and *K. pneumonia* using a final source concentration of 40, 10, 20 mg/L, respectively, while gentamicin, ofloxacin and tetracycline were tested against *S. aureus* using a final concentration of 2.5, 2.5 and 5 mg/L, respectively. Prior to the start of an experiment, the antibiotics were thawed and diluted in growth medium (MH-II broth, Becton Dickinson).

### Antibiotic susceptibility testing

Both the growth medium (MH-II broth) and the fluidic chip were degassed for 1 hour prior to every experiment to minimize the risk for formation of gas bubbles within the fluidic system. Suspensions of bacteria were prepared by dissolving 2-4 colonies in physiological NaCl where after the suspension was adjusted to 0.5 McFarland. Next, the suspension was further diluted 1:50 in growth medium and mixed 1:1 with 1% agarose (TopVision Low Melting Point Agarose, Fisher Scientific). Each chamber in the fluidic chip was loaded with 5 μL bacteria-agarose and incubated at 4 °C for 5 min to solidify the agarose gel. The chip was thereafter connected to the reservoir lid and assembled with the bottom plate and the top plates using M3 screws. Finally, tubing was inserted into the two fluid outlets on the chip and connected to two separate syringes fitted in a syringe pump set to produce a flow of 1 μL/min in the fluidic channels on each side of the growth chamber. The chip was primed with growth medium prior to every experiment and excess liquid removed from the lid reservoirs before growth medium with or without antibiotics was added to the appropriate reservoirs. The fully assembled chip was placed in a custom-built darkfield microscope fitted with a motorized camera module and a circuit board with heat elements. All experiments were carried out at 37°C. Images of each growth chamber were taken every 10 minutes using a Basler ACE camera (Basler ace acA2500-14uc Color, Basler). Data on BMD testing was obtained from the EUCAST Development Laboratory (EDL) in Växjö, Sweden.

### Image and data analysis

Image data was analysed in Python 2.7.10 using python libraries scipy (version 0.16.1), numpy (version 1.9.3), pandas (version 0.17.0), matplotlib (version 1.5.0rc3) and skimage (version 0.11.3). Image stacks were sorted based on chamber and read as 8-bit images (1942 x 2590 pixels). Positioning deviations were corrected using convolution and the images were cropped to only contain the actual chamber. To reduce background noise, a Gaussian filter was applied to the first image in the stack and the resultant background image was subtracted from the following images in the stack. Binary images were made using a threshold value of 10 units. Starting from the fifth image, the area of each microcolony was measured and compared to the previous 4 images in the stack. The growth rate of each microcolony was computed by fitting data to a linear function, where the slope indicates the rate of growth. Growth rates ≤0 or >100 were likely due to noise and therefore excluded. The chamber was divided into 500 equally spaced bins (approximately 5 px/bin) and the microcolony regions of interest were divided into different bins based on their x-coordinate. The mean of each bin was computed and data was fitted into a Gaussian distribution. The MIC was read at the first position where the Gaussian distribution < 1. After the MIC-signal over time had stabilized, and 3 values in a row were within 5% variation, the MIC was considered stable and was reported. Accordingly, the limit of quantification (LOQ) was defined as 1/20 of the maximum antibiotic concentration. For chambers with no signal, a read-out was performed 30 minutes after growth was detected in the positive growth control chamber. No MIC values were accepted before growth had been detected in the positive growth control chamber.

For comparison of essential agreement with reference BMD MICs, the obtained MIC values were right-censored to the nearest 2-log value, to allow for comparison on the same scale. As per the method used for previous method comparisons of methods with different concentration ranges (20), when the reference method showed results below or above LOQ for the comparative method, these were counted as in agreement. For comparison of categorical agreement, the MIC values from the essential agreement analysis were categorized by applying the EUCAST breakpoints for susceptible (S), sensitive with Increased exposure (I) and resistant (R) categories (The European Committee on Antimicrobial Susceptibility Testing, breakpoint tables for interpretation of MICs and zone diameters, version 9.0, 2019, see http://www.eucast.org/clinical_breakpoints/). When compared to BMD, S instead of R was counted as very major error (VME), R instead of S as major error (ME) and S or R instead of I as minor error (MiE).

## Author contributions (CRedIT categories)

Conceptualization: CM PWY TT JK

Data curation: PWY CM NFK

Formal analysis: CM PWY

Funding acquisition: TT JK

Investigation: PWY NFK CM

Methodology: PWY NFK CM

Project administration: CM PWY ML

Resources: TT JK

Software: PWY

Supervision: TT JK

Validation: CM PWY NFK ML

Visualization: PWY, CM, NFK

Writing – original draft: CM

Writing – review & editing: CM PWY NFK TT JK

## Acknowledgements

The authors wish to thank Gunnar Kahlmeter and Erika Matuschek at the EUCAST Development Laboratory for fruitful discussions, test strains and BMD susceptibility data. This study was funded by a grant to JK from Uppsala Antibiotic Centre, a grant to Gradientech AB and TT from Vinnova (grant nr. 2016-02286), and by grant to Gradientech and JK from the European Union’s Horizon 2020 research and innovation program under the Marie Sklodowska-Curie grant agreement 642866.

## References

1. Hughes D, Karlén A. 2014. Discovery and preclinical development of new antibiotics. Ups J Med Sci 119:162–169.

2. Moran GJ, Krishnadasan A, Gorwitz RJ, Fosheim GE, McDougal LK, Carey RB, Talan DA. 2006. Methicillin-Resistant S. aureus Infections among Patients in the Emergency Department. New England Journal of Medicine 355:666–674.

3. Shorr AF. 2007. Epidemiology of Staphylococcal Resistance. Clin Infect Dis 45:S171–S176.

4. Goossens H, Ferech M, Vander Stichele R, Elseviers M. 2005. Outpatient antibiotic use in Europe and association with resistance: a cross-national database study. The Lancet 365:579–587.

5. Auta A, Hadi MA, Oga E, Adewuyi EO, Abdu-Aguye SN, Adeloye D, Strickland-Hodge B, Morgan DJ. 2019. Global access to antibiotics without prescription in community pharmacies: A systematic review and meta-analysis. Journal of Infection 78:8–18.

6. Hughes JS, Hurford A, Finley RL, Patrick DM, Wu J, Morris AM. 2016. How to measure the impacts of antibiotic resistance and antibiotic development on empiric therapy: new composite indices. BMJ Open 6:e012040.

7. Spyropoulou A, Papadimitriou-Olivgeris M, Bartzavali C, Vamvakopoulou S, Marangos M, Spiliopoulou I, Anastassiou ED, Christofidou M. 2016. A ten-year surveillance study of carbapenemase-producing Klebsiella pneumoniae in a tertiary care Greek university hospital: predominance of KPC- over VIM- or NDM-producing isolates. Journal of Medical Microbiology 65:240–246.

8. Magiorakos A-P, Suetens C, Monnet DL, Gagliotti C, Heuer OE. 2013. The rise of carbapenem resistance in Europe: just the tip of the iceberg? Antimicrob Resist Infect Control 2:6.

9. Belkum A van, Bachmann TT, Lüdke G, Lisby JG, Kahlmeter G, Mohess A, Becker K, Hays JP, Woodford N, Mitsakakis K, Moran-Gilad J, Vila J, Peter H, Rex JH, Dunne WM. 2019. Developmental roadmap for antimicrobial susceptibility testing systems. Nature Reviews Microbiology 17:51.

10. Zhang D, Micek ST, Kollef MH. 2015. Time to Appropriate Antibiotic Therapy Is an Independent Determinant of Postinfection ICU and Hospital Lengths of Stay in Patients With Sepsis*: Critical Care Medicine 43:2133–2140.

11. Kumar A, Roberts D, Wood KE, Light B, Parrillo JE, Sharma S, Suppes R, Feinstein D, Zanotti S, Taiberg L, Gurka D, Kumar A, Cheang M. 2006. Duration of hypotension before initiation of effective antimicrobial therapy is the critical determinant of survival in human septic shock. Crit Care Med 34:1589–1596.

12. Dubourg G, Lamy B, Ruimy R. 2018. Rapid phenotypic methods to improve the diagnosis of bacterial bloodstream infections: meeting the challenge to reduce the time to result. Clinical Microbiology and Infection 24:935–943.

13. Chandrasekaran S, Abbott A, Campeau S, Zimmer BL, Weinstein M, Thrupp L, Hejna J, Walker L, Ammann T, Kirn T, Patel R, Humphries RM. 2018. Direct-from-Blood-Culture Disk Diffusion To Determine Antimicrobial Susceptibility of Gram-Negative Bacteria: Preliminary Report from the Clinical and Laboratory Standards Institute Methods Development and Standardization Working Group. Journal of Clinical Microbiology 56:e01678–17.

14. Pulido MR, García-Quintanilla M, Martín-Peña R, Cisneros JM, McConnell MJ. 2013. Progress on the development of rapid methods for antimicrobial susceptibility testing. J Antimicrob Chemother 68:2710–2717.

15. Moradigaravand D, Palm M, Farewell A, Mustonen V, Warringer J, Parts L. 2018. Prediction of antibiotic resistance in Escherichia coli from large-scale pan-genome data. PLOS Computational Biology 14:e1006258.

16. Khan ZA, Siddiqui MF, Park S. 2019. Progress in antibiotic susceptibility tests: a comparative review with special emphasis on microfluidic methods. Biotechnol Lett 41:221–230.

17. Malmberg C, Yuen P, Spaak J, Cars O, Tängdén T, Lagerbäck P. 2016. A Novel Microfluidic Assay for Rapid Phenotypic Antibiotic Susceptibility Testing of Bacteria Detected in Clinical Blood Cultures. PLOS ONE 11:e0167356.

18. Cho H, Uehara T, Bernhardt TG. 2014. Beta-Lactam Antibiotics Induce a Lethal Malfunctioning of the Bacterial Cell Wall Synthesis Machinery. Cell 159:1300–1311.

19. Buijs J, Dofferhoff ASM, Mouton JW, van der Meer JWM. 2007. Continuous administration of PBP-2-and PBP-3-specific β-lactams causes higher cytokine responses in murine Pseudomonas aeruginosa and Escherichia coli sepsis. J Antimicrob Chemother 59:926–933.

20. Reynolds R, Shackcloth J, Felmingham D, MacGowan A. 2003. Comparison of BSAC agar dilution and NCCLS broth microdilution MIC methods for in vitro susceptibility testing of Streptococcus pneumoniae, Haemophilus influenzae and Moraxella catarrhalis: the BSAC Respiratory Resistance Surveillance Programme. J Antimicrob Chemother 52:925–930.

21. Lat A, Clock SA, Wu F, Whittier S, Della-Latta P, Fauntleroy K, Jenkins SG, Saiman L, Kubin CJ. 2011. Comparison of Polymyxin B, Tigecycline, Cefepime, and Meropenem MICs for KPC-Producing Klebsiella pneumoniae by Broth Microdilution, Vitek 2, and Etest. Journal of Clinical Microbiology 49:1795–1798.

22. Dai J, Hamon M, Jambovane S. 2016. Microfluidics for Antibiotic Susceptibility and Toxicity Testing. Bioengineering 3:25.

23. Dong T, Zhao X. 2015. Rapid identification and susceptibility testing of uropathogenic microbes via immunosorbent ATP-bioluminescence assay on a microfluidic simulator for antibiotic therapy. Anal Chem 87:2410–2418.

24. Lu Y, Gao J, Zhang DD, Gau V, Liao JC, Wong PK. 2013. Single Cell Antimicrobial Susceptibility Testing by Confined Microchannels and Electrokinetic Loading. Anal Chem 85:3971–3976.

25. Baltekin Ö, Boucharin A, Tano E, Andersson DI, Elf J. 2017. Antibiotic susceptibility testing in less than 30 min using direct single-cell imaging. PNAS 114:9170–9175.

26. Haubert K, Drier T, Beebe D. 2006. PDMS bonding by means of a portable, low-cost corona system. Lab on a Chip 6:1548–1549.

