## Supplemental Figures for "A high-throughput fluidic chip for rapid phenotypic antibiotic susceptibility testing"

### Supplementary information

**Supplementary Material S1.** Summary of the data for all analyzed strains. Growth controls (untreated) are shown to the left.

### Gram-negative strains

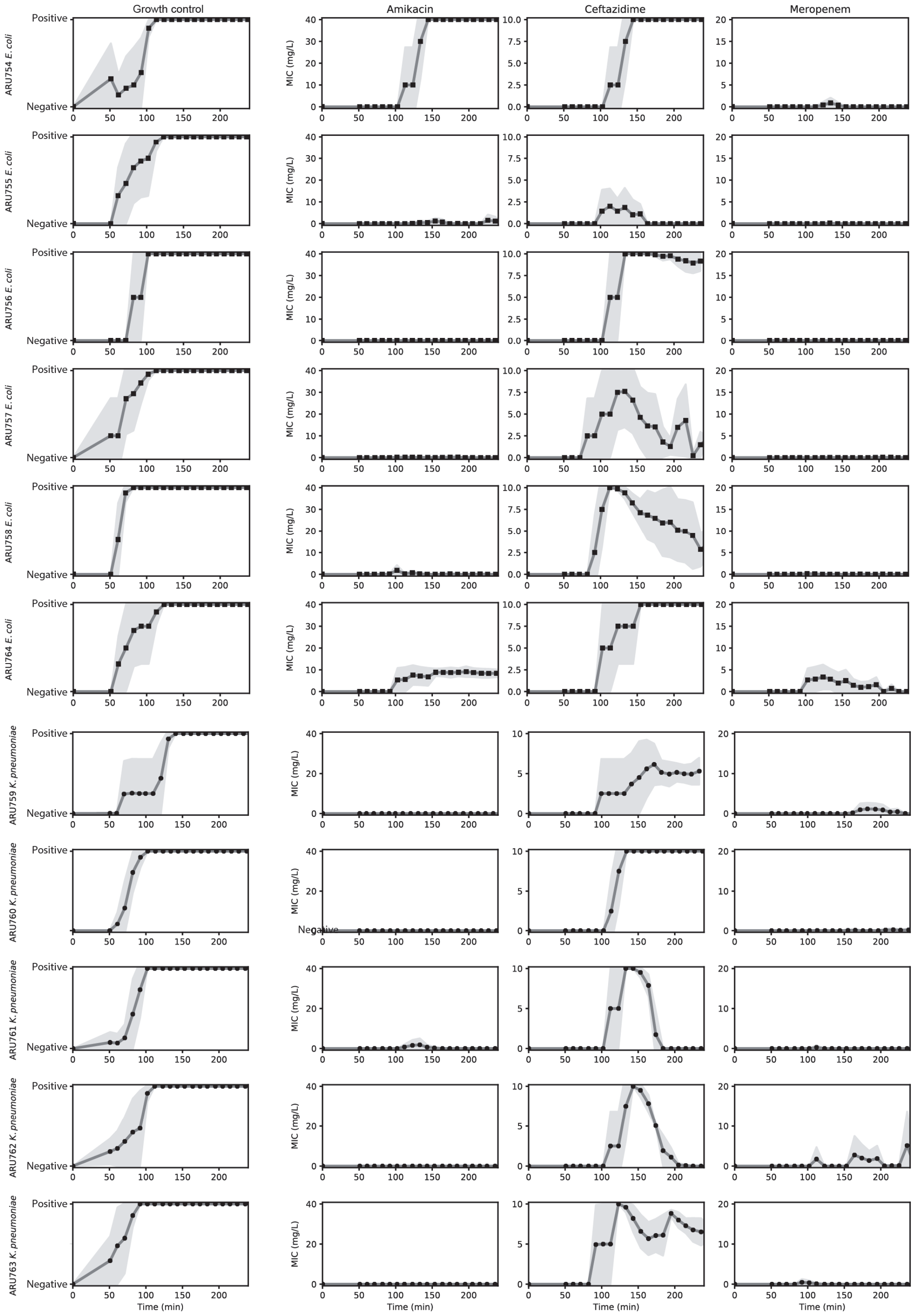


4

#### Gram-positive strains

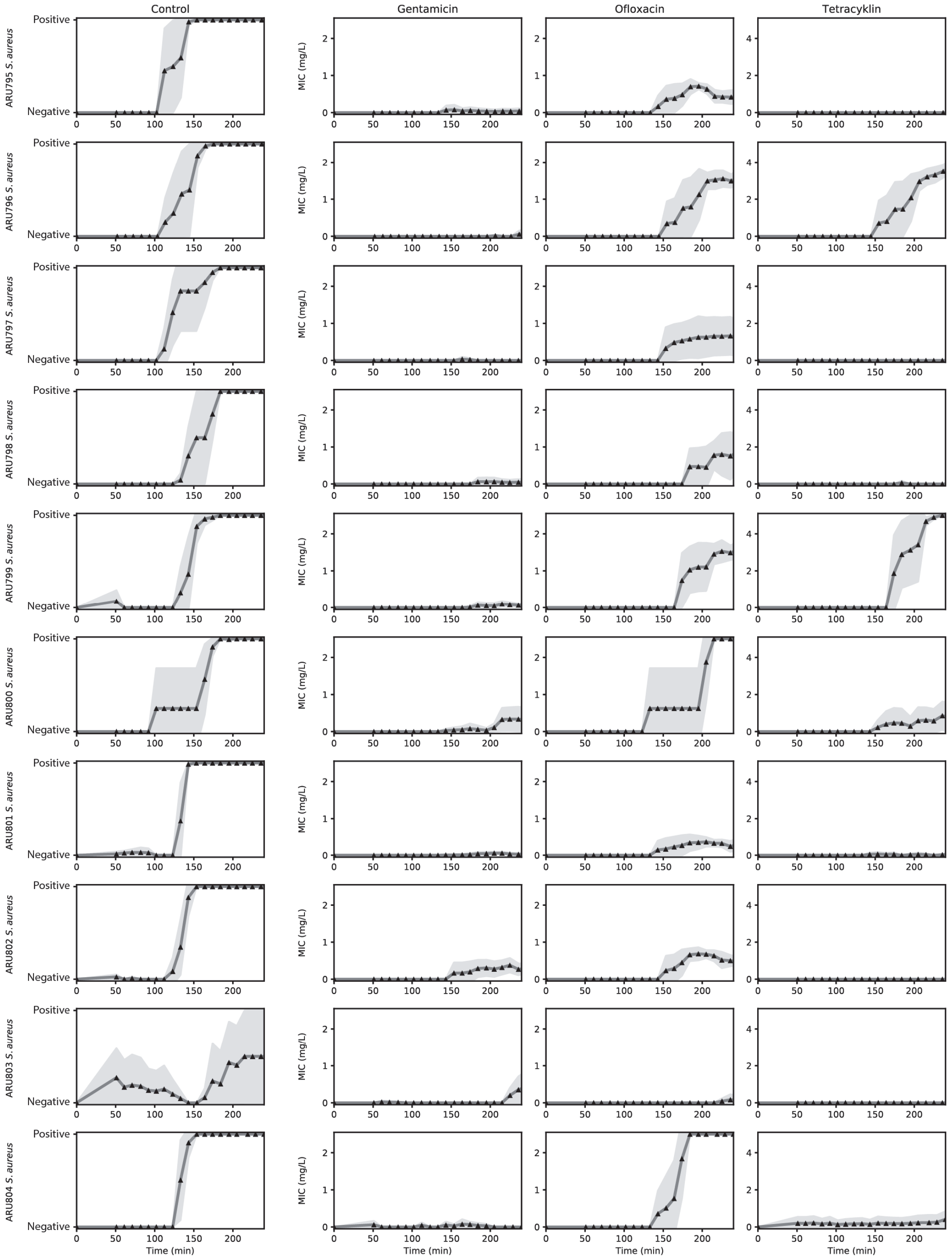
